# Multiscale Simulation of an Influenza A M2 Channel Mutant Reveals Key Features of Its Markedly Different Proton Transport Behavior

**DOI:** 10.1101/2021.09.07.459301

**Authors:** Laura C. Watkins, William F. DeGrado, Gregory A. Voth

## Abstract

The influenza A M2 channel, a prototype for the viroporin class of viral channels, is an acid-activated viroporin that conducts protons across the viral membrane, a critical step in the viral life cycle. As the protons enter from the viral exterior, four central His37 residues control the channel activation by binding subsequent protons, which opens the Trp41 gate and allows proton flux to the viral interior. Asp44 is essential for maintaining the Trp41 gate in a closed state at high pH, which results in asymmetric conduction. The prevalent D44N mutant disrupts this gate and opens the C-terminal end of the channel, resulting in overall increased conduction in the physiologically relevant pH range and a loss of this asymmetric conduction. Here, we use extensive Multiscale Reactive Molecular Dynamics (MS-RMD) and Quantum Mechanics/Molecular mechanics (QM/MM) simulations with an explicit, reactive excess proton to calculate the free energy of proton transport in the M2 mutant and to study the dynamic molecular-level behavior of D44N M2. We find that this mutation significantly lowers the barrier of His37 deprotonation in the activated state and shifts the barrier for entry up to the Val27 tetrad. These free energy changes are reflected in structural shifts. Additionally, we show that the increased hydration around the His37 tetrad diminishes the effect of the His37 charge on the channel’s water structure, facilitating proton transport and enabling activation from the viral interior. Altogether, this work provides key insight into the fundamental characteristics of PT in WT M2 and how the D44N mutation alters this PT mechanism, and it expands our understanding of the role of emergent mutations in viroporins.

## INTRODUCTION

Viroporins are small viral ion channels that are crucial for viral pathogenicity.^1^ They are found in many clinically relevant viruses including coronaviruses, influenza A virus, HIV-1, and hepatitis C virus, and are ideal therapeutic targets. The influenza A matrix protein 2 (M2) proton channel is considered a prototype for viroporins and is the target of the antiviral drugs amantadine and rimantadine.^2^ Following the encapsulation of the virus in a cellular endosome, M2 is responsible for the acidification of the viral interior via unidirectional proton transport (PT), which allows the virus to escape the endosome and infect the cell.^3–5^ The prevalent Asp44 to Asn (D44N) mutation results in increased proton conduction, which helps protect more acid-labile hemagglutinin in certain influenza strains.^6–8^ While much is known about PT in wildtype (WT) M2, the increased conduction in the D44N mutant is not as well understood. Studying this viroporin is important both for increasing our understanding of PT in M2, as well as for identifying the fundamental principles of PT through viroporins such as the SARS-CoV-2 ORF3a protein.^9^

M2 contains four central histidine residues (His37) that select for protons and control the channel’s activation.^10–12^ At high pH, all four residues are deprotonated, which we refer to as the +0 state, and the C-terminus of the channel is contracted in an inactive C_closed_ conformation. As the pH is lowered, each histidine residue can bind one additional proton, up to the +4 state. It is believed that most proton conduction occurs by cycling through the +2 and +3 states based on experimentally determined pK_a_’s, pH-dependent conduction, and multiscale computer simulations. ^11–17^ As a third excess proton enters the channel in a +2 state and binds to His37, electrostatic repulsion between these residues causes the C-terminal ends of the helices to widen into a C_open_ conformation which opens the Trp41 gate,^18–22^ increasing hydration and facilitating inward proton flux.^11, 12, 23–27^ NMR experiments of M2 at low pH show a broadening of peaks, suggesting a dynamic and flexible ensemble of C_open_ conformers.^22, 28^

One critical characteristic of PT in M2 is its asymmetric conductance, or inward rectification — WT M2 only becomes activated and conducts protons when the exterior pH is low.^29, 30^ When only the interior pH is lowered, M2 does not show any significant outward current, indicating that it cannot reach the necessary +2 charge state from internal protons. This asymmetry is moderated by the Trp41 gate, which forms a hydrophobic barrier in the C_closed_ state.^18^ With little water access to the His37 tetrad, the channel does not easily take up an excess proton from the interior and does not activate. Mutation of Trp41 removes this gate, resulting in outward conduction under low pH_in_ conditions.^18^ The D44N mutant also does not exhibit asymmetric conductance and can activate at low pH_in_. Previous studies have shown that Asp44 plays a particularly important structural role in stabilizing the closed conformation of the Trp41 gate.^22, 25, 29^ The four Asp carboxylates in the tetramer form an extensive network of hydrogen-bonds that include a salt bridge to Arg45 from neighboring helices, and a network of waters that hydrogen-bond to the Trp41 indole NH groups (**Figure 1**). The D44N mutation is expected to disrupt the salt bridge and the water-mediated hydrogen bonds to the Trp indole groups. Indeed, NMR studies by Ma, et al, show that this substitution destabilizes the Trp gate,^6^ increasing its dynamics and hydration. This resulting destabilized gate was hypothesized to allow unimpeded protonation and deprotonation of His37 from either side of the bilayer. These conclusions were supported by relatively short MD simulations, but the dynamics of the D44N mutant impeded attempts to structurally characterize D44N.

**Figure 1.**
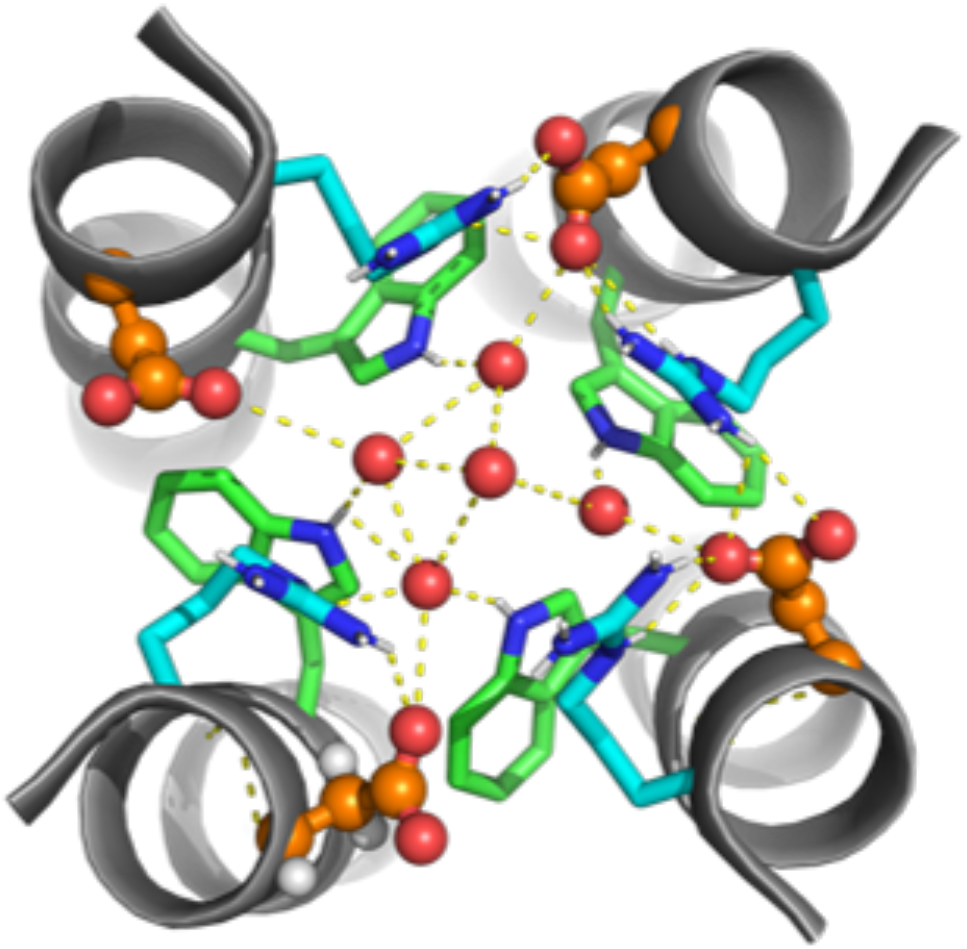
Structure of the Trp41 gate in WT M2 (PDB: 6US8^31^). The carbon atoms of Trp41, Asp44 and Arg 45 are, respectively in green, orange, and cyan. Water molecules are shown as red spheres.

Additional experiments by Ma, et al, confirmed that Asp44 is essential for asymmetric conductance.^6^ This work also showed that the D44N mutant has increased conduction and that the inward conduction has an altered pH_out_ dependence relative to WT. In the physiologically relevant pH range 6-5 (the pH of the endosome is approximately 6.0-5.4), the D44N mutant M2 is 3- to 4-fold more conductive than WT M2. Additionally, the pK_a_ values of two distinct protonation states relevant to conduction (hypothesized to be the formation of the +3 and +4 states) are shifted up in the D44N mutant, indicating that channel activation occurs at a higher pH. NMR and classical MD simulations in that study also showed that the D44N mutation induces conformational changes similar to those in WT at low pH.

It has not been studied and quantified, however, how the D44N mutation and corresponding structural changes affect PT on the molecular level, including the explicit proton transport. To this end, we have utilized extensive Multiscale Reactive Molecular Dynamics (MS-RMD)^32–35^ and Quantum Mechanics/Molecular Mechanics (QM/MM) simulations with an explicitly transporting excess proton to study PT in the transmembrane portion of D44N M2. These simulation methods allow one to address dynamical questions that are not accessible using classical (non-reactive) MD, which does not allow the bond-breaking and bond-making steps involved in proton transfer. MS-RMD was developed to efficiently and accurately model the solvation and delocalization of a hydrated excess proton in water within a classical MD context for aqueous and biomolecular systems. With potentials fit from accurate QM calculations,^36^ MS-RMD allows protons to form and break bonds with water molecules, capturing the underlying physics of the Grotthuss shuttling (proton hopping) mechanism,^37–39^ while also reaching the many-nanosecond timescales necessary for studying PT in complex biomolecular systems. MS-RMD has successfully been applied to several protein systems to predict and explain PT mechanisms, including influenza A M2.^11, 12, 40–49^

In our previous work,^11, 12^ MS-RMD and QM/MM simulations of the WT M2 channel were used to calculate potentials of mean force (PMFs) of explicit PT in the +0 through +3 protonation states, providing fundamental insight into the PT mechanism and role of the His37 tetrad. Recently, we further used these simulations to explore in detail how the hydrated excess proton interacts with the channel, and found that the excess proton dynamically, as a function of position, alters the protein structure and water hydrogen-bonding network.^46^ This approach was subsequently used to examine the precise interactions between the excess proton, water, and channel in the adamantine binding pocket.^49^ There, we showed that the channel is especially stable in the binding region, and we demonstrated how amantadine inhibitors take advantage of the channel’s natural ability to stabilize an excess proton charge.

Here, we employ MS-RMD and QM/MM simulations to deduce how the PT mechanism in D44N M2 differs from WT, how the mutation results in increased conductance, and why rectification is lost. The MS-RMD method is used to simulate PT in the +0, +1, and +2 protonation states through the channel. QM/MM simulations were used for PT through the His37 region, and together with MS-RMD were used to calculate the PMF of PT through the +2 state. Our results show that the +0 and +1 mutant states are structurally similar to the WT +2 state, and the +2 mutant state is significantly more open at the C-terminal. This opening causes the N-terminal end to be more closed, shifting the PMF barrier of entry to the Val27 gate. Additionally, we show how the altered water structures in D44N M2 facilitate PT, making the overall barrier for PT lower and thus increasing conductance, and enabling activation at low pH_in_.

## METHODS

### Classical MD Simulations

Because no experimental structures of D44N M2 are available, the initial M2 mutant structure was obtained by taking a previously equilibrated WT M2 structure from ref^12^ and mutating the residue in Visual Molecular Dynamics (VMD).^50^ The WT starting structure was resolved at room temperature and high pH (PDB: 4QKL^51^), embedded in a 1-palmitoyl-2-oleoyl-*sn*-glycero-3-phosphocholine (POPC) bilayer and solvated with water. This structure contains the transmembrane portion of M2, which is the minimum construct necessary to retain proton conduction similar to full-length M2.^52^ The mutant structure was initialized in the +0 state and equilibrated for at least 500 ns. The +1 and +2 states were generated by subsequently protonating one His37 residue and equilibrating for 500 ns twice.

### MS-RMD Simulations

MS-RMD was used to model the water and excess proton in simulations used in the protein and water structure analyses. The MS-RMD method allows hydrogen-oxygen bonds to break and form by taking a linear combination of possible bonding topology states at each timestep; we refer the reader to previous work for a full description of the method.^32–35^ The MS-EVB 3.2 parameters were used to describe the hydrated excess proton.^36^ The excess proton center of excess charge (CEC) is defined as^53^

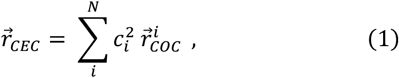

where 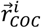 is the coordinate of the center of excess charge of the *i*th diabatic state and 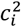 is amplitude of that state. The CEC allows one to track during the simulation the position of the most hydronium-like structure during the Grotthuss proton shuttling process through water molecules. The CHARMM36 force field was used to model the remaining interactions in the system. Simulations were run at 308 K in the NVT ensemble using LAMMPS^54^ with the MS-RMD package.

The collective variable (CV) used to define the excess proton’s progression through the channel was defined as the z-coordinate of the vector between the CEC and the center of mass of the four Gly34 α-carbons as in our previous work.^11, 12, 46, 49^ Umbrella sampling (US) was performed using the replica exchange umbrella sampling (REUS)^55^ technique to facilitate convergence. Simulations were run with the proton restrained every 0.5 Å along the CV coordinate throughout the channel, minus the His37 region, with a 10.0 kcal·mol^-1^·Å^-2^ US force constant. Additionally, a cylindrical restraint was applied to ensure the proton remained in the channel at 8.0 Å and 12.0 Å for the top and bottom of the channel, respectively, with a force constant of 10.0 kcal·mol^-1^·Å^-2^using the open-source, community developed PLUMED library.^56, 57^ Windows were equilibrated for 250 ps, followed by 1.5-5 ns production simulations.

### QM/MM Simulations

While MS-RMD has been successfully used to model amino acid residue protonation and deprotonation in certain previous work, there is not currently a reliable method for fitting models of coupled residues in very complex environments, such as the His37 tetrad. Thus, to calculate the PMF for PT through the M2 channel in the +2 state, QM/MM simulations were run for PT through the His37-Trp41 tetrad region following the same protocol as in our previous work.^11, 12^ The CEC coordinate used in the QM/MM simulations is fully defined in the Supporting Information. This formulation captures the delocalized nature of the excess proton; it has been shown to adequately describe the excess proton position in QM/MM simulations of biological PT channels^58–60^ and has been used in our previous work with M2.^11, 12^

Simulation parameters are the same as in our previous work,^12^ but briefly summarized here. The QM atoms included the His37 sidechains, the excess proton, and up to three solvation shells of water above and below His37 to ensure the proton was sufficiently surrounded by QM waters. The QM region was treated by Becke-Lee-Yang-Parr level density functional theory^61, 62^ with empirical dispersion corrections,^63^ under the Gaussian plane wave scheme.^64^ Goedecker-Teter-Hutter pseudopotentials^65^ were used and the Kohn-Sham orbitals were expanded in the Gaussian TZV2P basis set. The integration timestep was 0.5 fs, and the temperature was set at 308 K and controlled by a Nose-Hoover thermostat with a 0.1 ps relaxation time.

The QM/MM simulations were set up as follows. Three initial windows were set up from equilibrated MS-RMD simulations: two with the excess proton on either end of the region, and one with the excess proton bound to one of the His37 residues. Umbrella sampling windows were spaced every ~0.25 Å for a total of 43 windows, and the QM CEC was restrained with a 40 kcal·mol^-1^·Å^-2^ force constant. Simulation windows were pulled from adjacent equilibrated windows. Each window had an equilibration time of ~2ps and ~12-35ps of production sampling. The CEC position was collected every step. Because the +2 channel is more open, MM water would occasionally enter the QM region. To address this, waters were checked every ~3ps of simulation and appropriately reassigned to the QM or MM region if needed. Systems were equilibrated for 100fs following such an exchange before continuing the production run. All QM/MM simulations were performed using the CP2K package.^66^

### PMF Calculations

These simulation results were combined with MS-RMD simulation results to calculate the hybrid PMF using the discrete transition-based reweighting analysis method (dTRAM)^67^ as implemented in the PyEMMA python package.^68^ Error bars were estimated using a block-averaging analysis.

### Analysis

All analyses in this paper were performed using MS-RMD US trajectories to delineate changes as the excess proton moves through the channel. Analysis was done with respect to the principal axis of the channel, Z’, to account for the channel’s tilt.^46^ The CEC position along this axis is labeled CEC_Z’_.

The direction of hydrogen bonds in the water analysis was calculated as the cosine of the angle between the donor-acceptor vector and the Z’ axis. See our previous work for more detail.^46, 49^

## RESULTS AND DISCUSSION

### D44N Mutation Lowers His37 Deprotonation Barrier and Shifts Proton Entry Barrier

Hybrid MS-RMD and QM/MM simulations were used to calculate the +2 mutant PMF shown in **Figure 2**, as described in the Methods. This PMF is the free energy associated with an additional excess proton moving through the channel in a +2 charge state. In the MS-RMD simulations, where the excess proton is out of the His37 tetrad region, the +2 charge is fixed and two histidine residues on opposing helices are protonated. In the QM/MM region, all four histidine residues and surrounding water are treated in the QM regime, allowing the protons to delocalize and the excess proton to form and break bonds with His37 residues.

**Figure 2.**
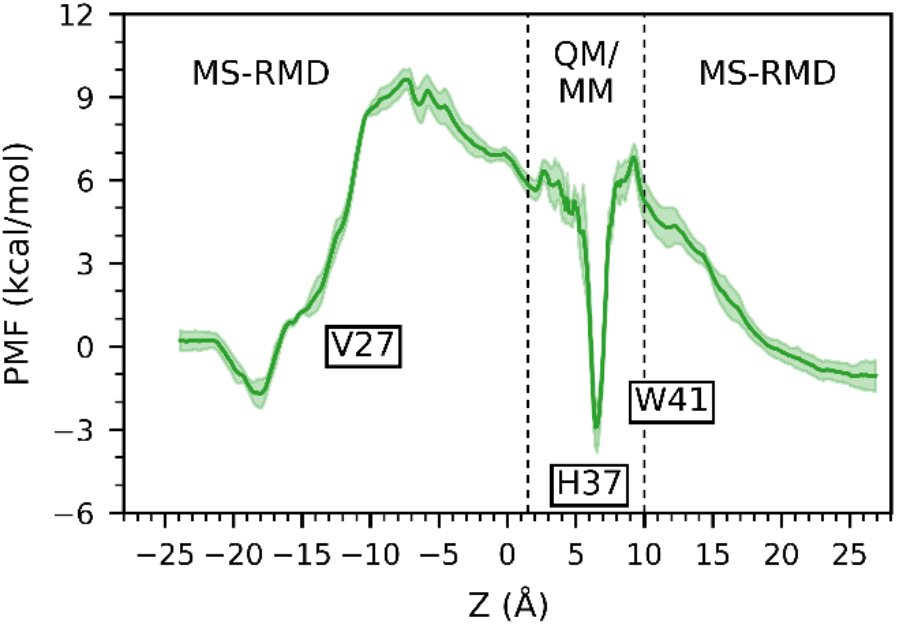
PMF for PT through the +2 D44N mutant state, calculated from hybrid MS-RMD and QM/MM simulations. The regions calculated from each simulation method are indicated. Error bars are shown.

There are two main features of the PMF that are distinct from the WT +2 PMF. First, the barrier for proton entry from the exterior is shifted from proton CEC position at Z = 0 Å in WT to Z = −10 Å in the mutant, at the location of Val27. This indicates that PT is more difficult through the Val27 tetrad, the primary gate at the top of the channel, relative to WT. Below, we show and discuss how the structural changes associated with the D44N mutation may make the top half more closed and rigid.

**Table 1.**
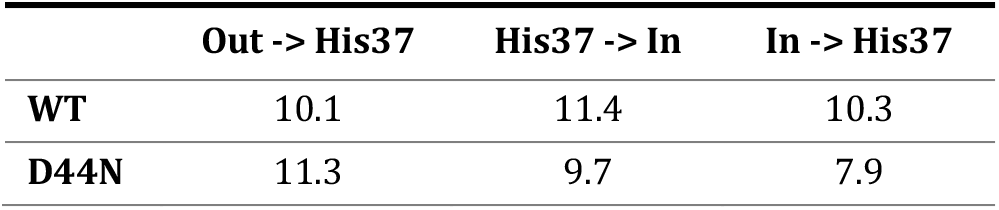
Barriers for different steps in the forward and reverse PT process from calculated +2 PMFs.^1^.

The second distinction from WT is a ~2.4 kcal/mol decrease in the barrier for proton entry from the interior and ~1.7 kcal/mol decrease for His37 deprotonation towards the viral interior, around Z = 9 Å. In WT M2, this deprotonation is the rate-limiting step in the PT process.^24, 69^ The lower barrier here indicates that protons can pass through the His37 tetrad more readily, resulting in overall increased conduction, in agreement with experiments. This decrease also changes which PT step has the largest free energy barrier—now, proton entry from the viral exterior becomes the largest barrier, indicating that in D44N M2 the rate-limiting step may be proton entry through the Val27 gate rather than His37 deprotonation.

### Structural Changes Due to D44N Mutation

The D44N mutation alters the protein’s structure and its equilibrium dynamics, causing the channel to be wider at the bottom compared to WT. To first examine the difference in structure, we compared the pore radii profiles between the WT and D44N mutant structures for the +0-+2 His37 charge states with no excess proton in the channel (**Figure 3**, top). In WT M2, with no excess proton in the channel, there is a narrow constriction at the Trp41 gate in the +0 and +1 structures. The +2 structure is more open, with a pore radius ~ 3Å, which allows water to freely reach the His37 tetrad and thus PT is facilitated. In the D44N mutant +0 and +1 states, the Trp41 gate is open with a pore radius of ~3Å, similar to the WT +2 state. The +2 mutant state is further expanded: below the Trp41 gate, the +2 state is much more open, with a 3Å increase in radius near the D44N mutation. Additionally, the +2 state is noticeably wider in the region between Gly34 and His37.

**Figure 3.**
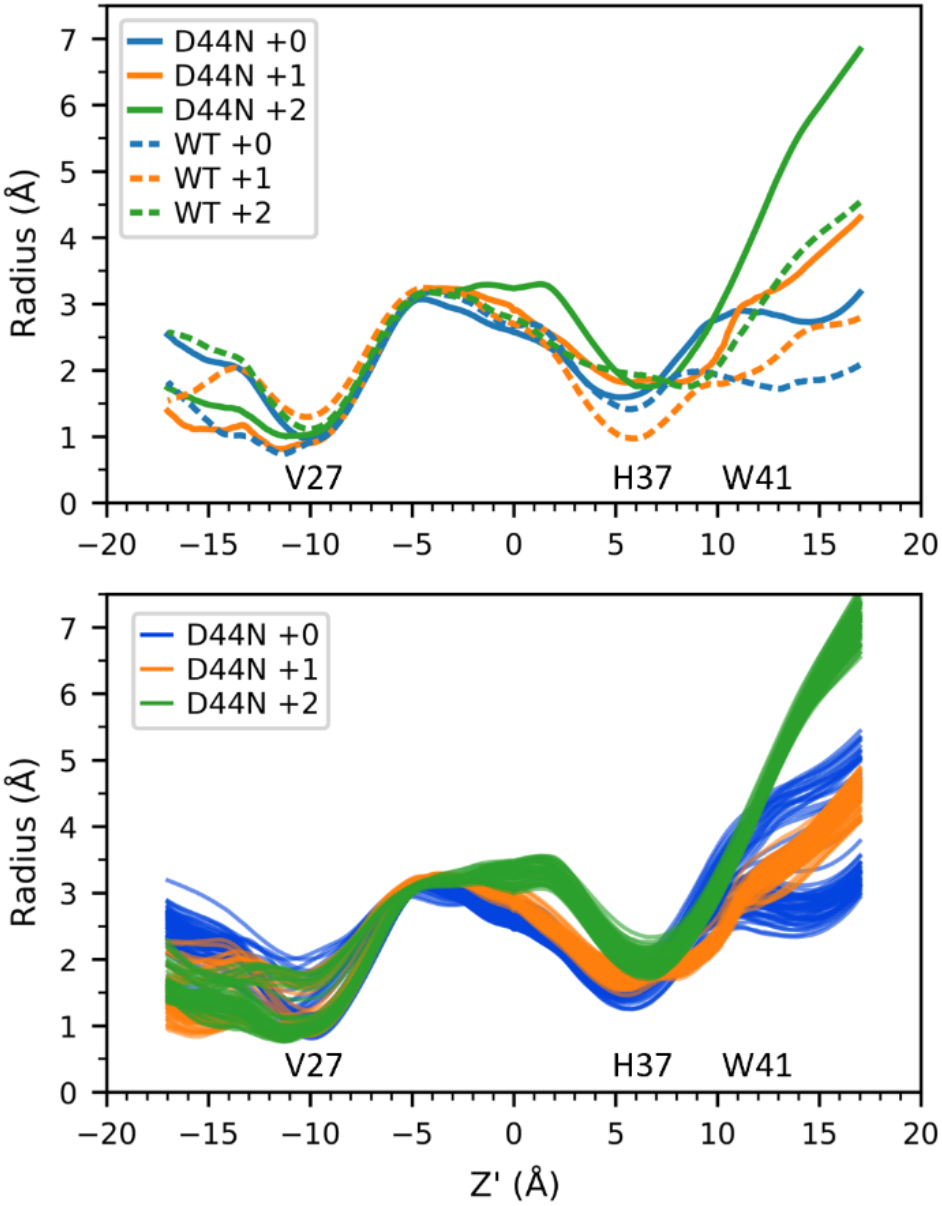
Top: radii profiles for WT and D44N M2 in the +0, +1, and +2 charge states. The x-axis is the Z’ channel axis. WT data is taken from ref. 43. These profiles represent the average structures when the excess proton is out of the channel (CEC_Z’_ = −24.0Å). Bottom: The average radii profiles calculated for excess proton positions every 0.25 Å in the mutant channel. Trajectory frames were binned by excess proton CEC_Z’_ with bin widths of 0.25Å. Each line is the average radii profile for a given bin.

Between Val27 and just above Gly34, Z’=-10.0 to Z’=-2.0 Å, the pore is remarkably similar to WT. We previously showed the stability in this region in WT M2 is a critical determinant for successful amantadine inhibition.^49^ Although the D44N mutation significantly alters the conformation of the channel, it surprisingly does not change this region. This finding is consistent with observation that the D44N M2 channel is sensitive to amantadine.^7^ In this case, drug binding induces the closed conformation with packed Trp41 indoles, although there are greater structural fluctuation in D44N versus WT near the His37/Trp41 gate. Above Val27, the +1 and +2 mutant structures are more closed relative to their WT counterparts.

We also looked at how the pore radii change as the proton moves through the channel by calculating the average channel radii profile for different excess proton positions throughout the channel (**Figure 3,** bottom). Each thin line in this figure shows the calculated radii profile when the excess proton CEC_Z’_ is fixed at a specific position. These profiles were calculated for CEC_Z’_ values every 0.25 Å between −22 Å and 22 Å (minus the QM/MM region), for a total of 144 lines for each charge state. Here, we see changes in the pore radii at the top of the channel in all three states, dependent on proton position, as well as at the bottom of the channel in the +0 state. In our previous work, we showed increased variation at His37 in the +0 and +1 WT structures that is not observed in the +2 state. This variation seen most prominently in +0 WT M2 occurs as the proton moves through the Val27 gate — for a proton to pass through the Val27 gate, +0 WT M2 transiently adopts a conformation that widens the top portion of the channel while constricting the bottom portion. We attributed this correlated movement between the top and bottom halves of the channel to an increased rigidity at high pH. In contrast, all three charge states of the mutant show minimal structural variation due to proton position near His37, suggesting that the C-terminal end of D44N M2 is more flexible than WT at high pH, similar to the low pH WT structures. There is, however, noticeable variation below W41.

The mutant’s shift towards a structure that is more open towards the viral interior and closed towards the viral exterior affects the pore hydration and free energy associated with PT — the free energy barrier for proton entry into the channel from the exterior increases, while the barrier for proton entry from the interior decreases, as reflected in the PMFs shown and discussed above.

### D44N Mutation Increases Hydration and Alters the Water Network

To understand how the structural differences in the D44N mutant affect the water structures and PT throughout the channel, we analyzed the number of waters within different regions of the channel and the water hydrogen bond orientations. In our previous work, we showed how the increased water and optimal orientation of the water near His37 in the WT +2 state facilitated PT and “primed” the +2 state for PT. Conversely, fewer waters and less optimal hydrogen-bonding orientation in the region below His37 in the +0 and +1 states with no excess proton present indicate that the water structure must change significantly for PT to occur through this region, as seen when the proton nears the His37 tetrad. The greater rearrangement necessary for PT from the viral interior to His37 in high pH structures also helps explain the rectification behavior of WT M2 — this rearrangement requirement increases the energy barrier for a proton to enter from the viral interior, and the water hydrogen bond network orientated away from the His37 tetrad suggests it would be easier for a proton to move off the tetrad than an additional proton to move in.

One critical component of this priming in the WT +2 state is the water structure between His37 and Trp41, which facilitates His37 deprotonation towards the viral interior. Throughout the PT process the +2 state consistently has 9-11 water molecules in this region, whereas the +0 and +1 states have fewer than 6, except when the proton is in this region and it draws an additional 2-3 waters in.^46^ In the mutant, all three charge states consistently have 8 or more waters in this region, with lower variability between states (**Figure 4**). The number of waters only dips below 8 in the +0 state when CEC_Z’_ = −2.0 Å, at which point the His37-Trp41 region is less relevant. This difference from WT suggests that the likelihood of proton release from His37 to the virial interior at high pH is increased in D44N M2, as the high pH structures represented by +0 and +1 have similar water structures to the activated, low pH +2 state. This increased likelihood suggests more leaky conductance at high pH.

**Figure 4.**
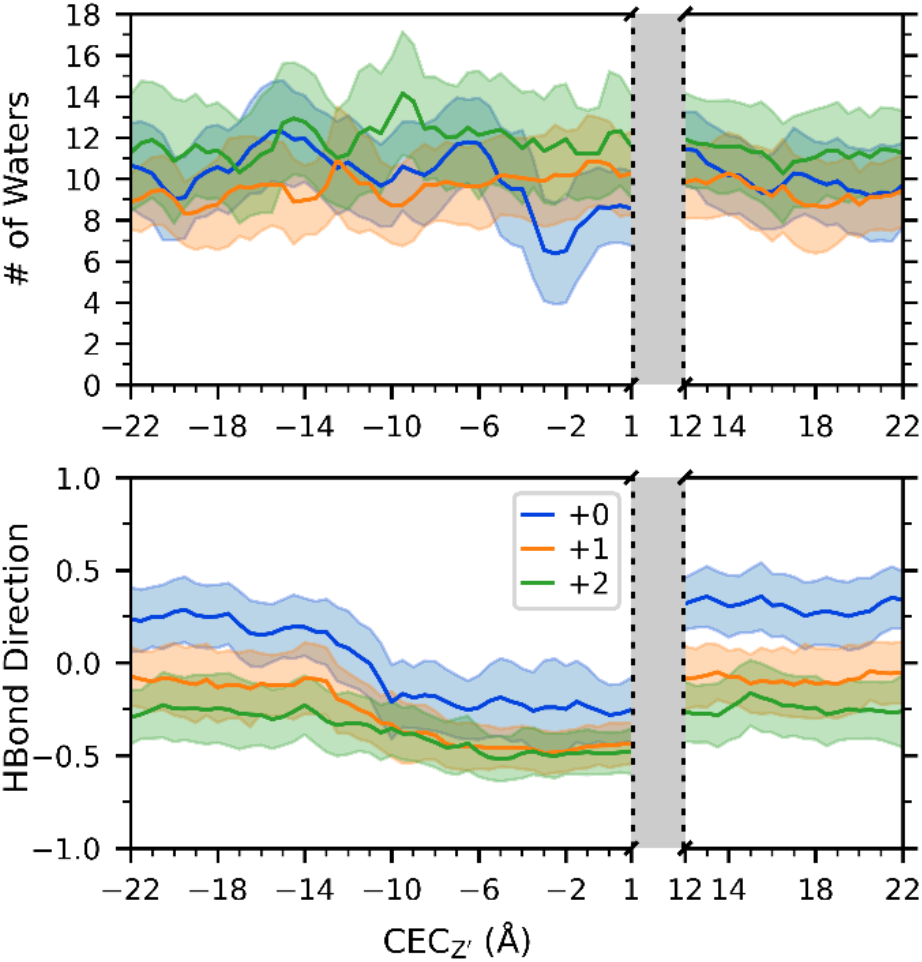
Water structure analysis for D44N in the region below His37 and above Trp41 from MS-RMD simulations. Top: The average and standard deviation of the number of waters in this region, as a function of the excess proton CEC position in the channel. Bottom: The average and standard deviation of the water-water hydrogen bonds direction. The grayed-out region corresponds to the QM/MM region.

The hydrogen bond network below the His37 tetrad in D44N is similar to that in WT, with the +0 state taking on overall positive average values with no proton present, and the +1 and +2 states have negative values. The +0 state value in D44N, however, is lower than in WT by ~0.25. This lowering suggests that the increase in number of waters limits the His37’s influence on water direction, which correlates with previous reports of a “parallel circuit” of hydrogen bonds in D44N M2 from classical MD simulations.^51^ This effect is most clearly seen in the region above Gly34 and below Ser31, as shown in **Figure 5**. In this region in WT, each state has a distinct line with values ~0.12, 0.23, 0.43 when an excess proton is not in the channel. In the mutant, all three charge states have similar values throughout the PT process, indicating that the His37 tetrad charge has minimal effect on the water structure in this region. The greater similarity in water networks across charge states in D44N M2 indicates that there may be fewer differences in PT between states in this mutant.

**Figure 5.**
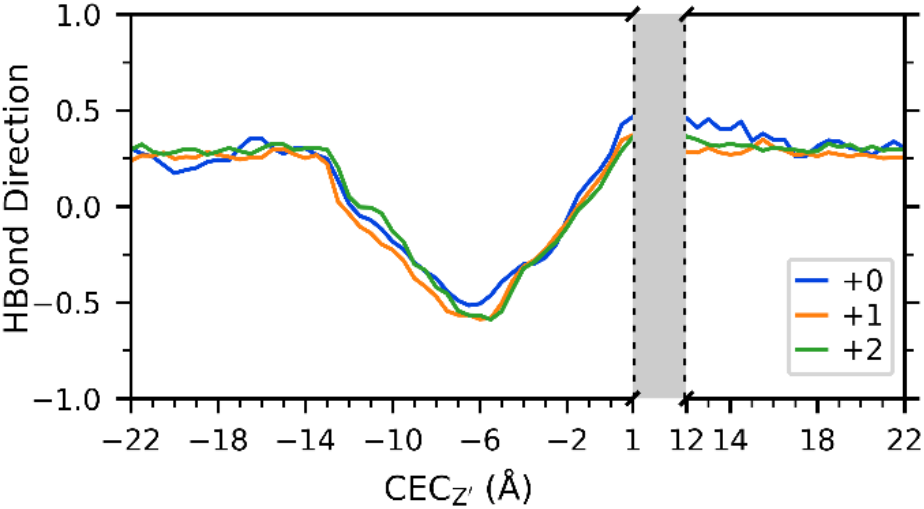
Average water-water hydrogen bond direction for D44N in the region above Gly34 and below Ser31 as a function of the excess proton CEC position in the channel. The grayed-out region corresponds to the QM/MM region.

The different water structure in D44N M2 also helps explain the loss of rectification behavior in this mutant, that is, that D44N M2 can become activated by protons entering from the viral interior at low pH_in_ and high pH_out_. In WT M2, the +0 and +1 states require additional water molecules below His37 for an excess proton to move through that region, whereas there is no significant change in water number in the +0 and +1 mutant states. Because the region is sufficiently hydrated, protons can more easily enter from the viral interior and the rate of protonation (*k_on_*) can compete with the rate of deprotonation towards the exterior (*k_off_*). If *k_on_* outcompetes *k_off_*, it becomes significantly probable to reach an activated +2 state. This is also reflected in the PMFs, as discussed above, which show that the barrier for protonation from the interior is ~2 kcal/mol less in the mutant M2 than in WT M2.

## CONCLUSIONS

In this work, we have used extensive MS-RMD and QM/MM simulations with an explicit excess proton to investigate how PT in the D44N M2 mutant differs from WT on the molecular level. First, we calculated the PMF for PT through the +2 state and discussed critical differences from WT—the barrier for His37 protonation from the viral interior is reduced, and the barrier for entry from the viral exterior is shifted to the Val27 gate, increasing conduction and potentially changing the rate-limiting step. We furthered our understanding of how this mutation alters the structure of M2 by examining channel radii profiles, showing that the high pH structures are similar to the low pH WT structure, and that opening the bottom of the channel results in narrowing the pore at the top near Val27. Finally, by analyzing the water hydrogen bond network, we showed that increased hydration facilitates PT and His37 binding in the low charge states and decreases the effect of the His37 charge on the water orientation. Together, these results also help explain the loss of rectification in the D44N mutant.

Studying the prevalent mutants of M2 is critical for continuing anti-flu drug-design efforts and is necessary to fully understand PT in M2 and the function of specific residues. Here, we show the significance of Asp44 and its naturally occurring variants in determining fundamental characteristics of PT, including the level of conduction, rectification, and pH dependence, and how this mutation alters these characteristics. As a prototype for viroporins more generally, these results indicate the importance of investigating mutations to deduce their effect on transport mechanisms and their physiological relevance.

## ASSOCIATED CONTENT

### Supporting Information

A full description of the CEC definition used in QM/MM simulations is included in the SI. This material is available free of charge via the Internet at http://pubs.acs.org.

## AUTHOR INFORMATION

### Author Contributions

All authors have given approval to the final version of the manuscript.

## ACKNOWLEDGMENT

The personnel in this research were supported by the National Institute of General Medical Sciences (NIGMS) of the National Institutes of Health (NIH Grants R01 GM053148 to G.A.V and R35 GM122603 to W.F. D.). The authors acknowledge The University of Chicago Research Computing Center and the U.S. Department of Defense High Performance Computing Modernization Program for providing computing resources. L.C.W. received partial funding from a Department of Energy (DOE) Computational Science Graduate Fellowship under Grant DE-FG02-97ER25308.

## Insert Table of Contents artwork here

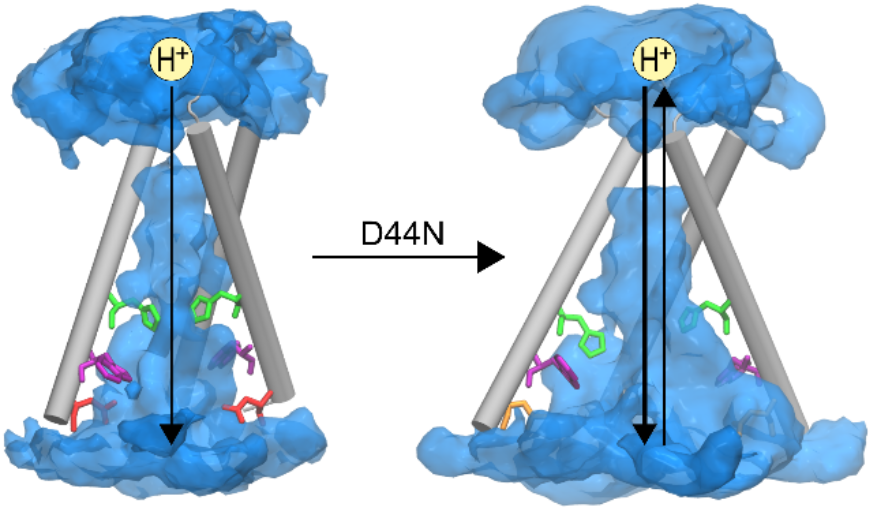

## Footnotes

1 All numbers are in kcal/mol. The three columns from left to right are for 1) proton entry from the viral exterior to His37 protonation, 2) His37 deprotonation towards the viral interior, and 3) proton entry from the viral interior to His37 protonation.

## REFERENCES

1. Nieva, J. L.; Madan, V.; Carrasco, L., Viroporins: structure and biological functions. Nat Rev Microbiol 2012, 10 (8), 563–74. DOI: 10.1038/nrmicro2820

2. Wang, C.; Takeuchi, K.; Pinto, L. H.; Lamb, R. A., Ion channel activity of influenza A virus M2 protein: characterization of the amantadine block. J Virol 1993, 67 (9), 5585–94. DOI: 10.1128/JVI.67.9.5585-5594.1993

3. Pinto, L. H.; Lamb, R. A., The M2 proton channels of influenza A and B viruses. J Biol Chem 2006, 281 (14), 8997–9000. DOI: 10.1074/jbc.R500020200

4. Chizhmakov, I. V.; Geraghty, F. M.; Ogden, D. C.; Hayhurst, A.; Antoniou, M.; Hay, A. J., Selective proton permeability and pH regulation of the influenza virus M2 channel expressed in mouse erythroleukaemia cells. J Physiol 1996, 494 (Pt 2) (2), 329–36. DOI: 10.1113/jphysiol.1996.sp021495

5. Pinto, L. H.; Holsinger, L. J.; Lamb, R. A., Influenza virus M2 protein has ion channel activity. Cell 1992, 69 (3), 517–528. DOI: 10.1016/0092-8674(92)90452-i

6. Ma, C.; Fiorin, G.; Carnevale, V.; Wang, J.; Lamb, R. A.; Klein, M. L.; Wu, Y.; Pinto, L. H.; DeGrado, W. F., Asp44 stabilizes the Trp41 gate of the M2 proton channel of influenza A virus. Structure 2013, 21 (11), 2033–41. DOI: 10.1016/j.str.2013.08.029

7. Betakova, T.; Ciampor, F.; Hay, A. J., Influence of residue 44 on the activity of the M2 proton channel of influenza A virus. J Gen Virol 2005, 86 (Pt 1), 181–184. DOI: 10.1099/vir.0.80358-0

8. Grambas, S.; Bennett, M. S.; Hay, A. J., Influence of amantadine resistance mutations on the pH regulatory function of the M2 protein of influenza A viruses. Virology 1992, 191 (2), 541–9. DOI: 10.1016/0042-6822(92)90229-i

9. Kern, D. M.; Sorum, B.; Mali, S. S.; Hoel, C. M.; Sridharan, S.; Remis, J. P.; Toso, D. B.; Kotecha, A.; Bautista, D. M.; Brohawn, S. G., Cryo-EM structure of SARS-CoV-2 ORF3a in lipid nanodiscs. Nat Struct Mol Biol 2021, 28 (7), 573–582. DOI: 10.1038/s41594-021-00619-0

10. Wang, C.; Lamb, R. A.; Pinto, L. H., Activation of the M2 ion channel of influenza virus: a role for the transmembrane domain histidine residue. Biophys J 1995, 69 (4), 1363–71. DOI: 10.1016/S0006-3495(95)80003-2

11. Liang, R.; Li, H.; Swanson, J. M.; Voth, G. A., Multiscale simulation reveals a multifaceted mechanism of proton permeation through the influenza A M2 proton channel. Proc Natl Acad Sci U S A 2014, 111 (26), 9396–401. DOI: 10.1073/pnas.1401997111

12. Liang, R.; Swanson, J. M. J.; Madsen, J. J.; Hong, M.; DeGrado, W. F.; Voth, G. A., Acid activation mechanism of the influenza A M2 proton channel. Proc Natl Acad Sci U S A 2016, 113 (45), E6955–E6964. DOI: 10.1073/pnas.1615471113

13. Miao, Y.; Fu, R.; Zhou, H. X.; Cross, T. A., Dynamic Short Hydrogen Bonds in Histidine Tetrad of Full-Length M2 Proton Channel Reveal Tetrameric Structural Heterogeneity and Functional Mechanism. Structure 2015, 23 (12), 2300–2308. DOI: 10.1016/j.str.2015.09.011

14. Liao, S. Y.; Yang, Y.; Tietze, D.; Hong, M., The influenza m2 cytoplasmic tail changes the proton-exchange equilibria and the backbone conformation of the transmembrane histidine residue to facilitate proton conduction. J Am Chem Soc 2015, 137 (18), 6067–77. DOI: 10.1021/jacs.5b02510

15. Colvin, M. T.; Andreas, L. B.; Chou, J. J.; Griffin, R. G., Proton association constants of His 37 in the Influenza-A M218-60 dimer-of-dimers. Biochemistry 2014, 53 (38), 5987–94. DOI: 10.1021/bi5005393

16. Hu, J.; Fu, R.; Nishimura, K.; Zhang, L.; Zhou, H. X.; Busath, D. D.; Vijayvergiya, V.; Cross, T. A., Histidines, heart of the hydrogen ion channel from influenza A virus: toward an understanding of conductance and proton selectivity. Proc Natl Acad Sci U S A 2006, 103 (18), 6865–70. DOI: 10.1073/pnas.0601944103

17. Hong, M.; DeGrado, W. F., Structural basis for proton conduction and inhibition by the influenza M2 protein. Protein Sci 2012, 21 (11), 1620–33. DOI: 10.1002/pro.2158

18. Tang, Y.; Zaitseva, F.; Lamb, R. A.; Pinto, L. H., The gate of the influenza virus M2 proton channel is formed by a single tryptophan residue. J Biol Chem 2002, 277 (42), 39880–6. DOI: 10.1074/jbc.M206582200

19. Hu, F.; Luo, W.; Cady, S. D.; Hong, M., Conformational plasticity of the influenza A M2 transmembrane helix in lipid bilayers under varying pH, drug binding, and membrane thickness. Biochim Biophys Acta 2011, 1808 (1), 415–23. DOI: 10.1016/j.bbamem.2010.09.014

20. Hu, F.; Luo, W.; Hong, M., Mechanisms of proton conduction and gating in influenza M2 proton channels from solid-state NMR. Science 2010, 330 (6003), 505–8. DOI: 10.1126/science.1191714

21. Li, C.; Qin, H.; Gao, F. P.; Cross, T. A., Solid-state NMR characterization of conformational plasticity within the transmembrane domain of the influenza A M2 proton channel. Biochim Biophys Acta 2007, 1768 (12), 3162–70. DOI: 10.1016/j.bbamem.2007.08.025

22. Schnell, J. R.; Chou, J. J., Structure and mechanism of the M2 proton channel of influenza A virus. Nature 2008, 451 (7178), 591–5. DOI: 10.1038/nature06531

23. Sharma, M.; Yi, M.; Dong, H.; Qin, H.; Peterson, E.; Busath, D. D.; Zhou, H. X.; Cross, T. A., Insight into the mechanism of the influenza A proton channel from a structure in a lipid bilayer. Science 2010, 330 (6003), 509–12. DOI: 10.1126/science.1191750

24. Polishchuk, A. L.; Lear, J. D.; Ma, C.; Lamb, R. A.; Pinto, L. H.; Degrado, W. F., A pH-dependent conformational ensemble mediates proton transport through the influenza A/M2 protein. Biochemistry 2010, 49 (47), 10061–71. DOI: 10.1021/bi101229m

25. Acharya, R.; Carnevale, V.; Fiorin, G.; Levine, B. G.; Polishchuk, A. L.; Balannik, V.; Samish, I.; Lamb, R. A.; Pinto, L. H.; DeGrado, W. F.; Klein, M. L., Structure and mechanism of proton transport through the transmembrane tetrameric M2 protein bundle of the influenza A virus. Proc Natl Acad Sci U S A 2010, 107 (34), 15075–80. DOI: 10.1073/pnas.1007071107

26. Wei, C.; Pohorille, A., Activation and proton transport mechanism in influenza A M2 channel. Biophys J 2013, 105 (9), 2036–45. DOI: 10.1016/j.bpj.2013.08.030

27. Williams, J. K.; Zhang, Y.; Schmidt-Rohr, K.; Hong, M., pH-dependent conformation, dynamics, and aromatic interaction of the gating tryptophan residue of the influenza M2 proton channel from solid-state NMR. Biophys J 2013, 104 (8), 1698–708. DOI: 10.1016/j.bpj.2013.02.054

28. Hu, J.; Asbury, T.; Achuthan, S.; Li, C.; Bertram, R.; Quine, J. R.; Fu, R.; Cross, T. A., Backbone structure of the amantadine-blocked trans-membrane domain M2 proton channel from Influenza A virus. Biophys J 2007, 92 (12), 4335–43. DOI: 10.1529/biophysj.106.090183

29. Chizhmakov, I. V.; Ogden, D. C.; Geraghty, F. M.; Hayhurst, A.; Skinner, A.; Betakova, T.; Hay, A. J., Differences in conductance of M2 proton channels of two influenza viruses at low and high pH. J Physiol 2003, 546 (Pt 2), 427–38. DOI: 10.1113/jphysiol.2002.028910

30. Mould, J. A.; Li, H. C.; Dudlak, C. S.; Lear, J. D.; Pekosz, A.; Lamb, R. A.; Pinto, L. H., Mechanism for proton conduction of the M(2) ion channel of influenza A virus. J Biol Chem 2000, 275 (12), 8592–9. DOI: 10.1074/jbc.275.12.8592

31. Thomaston, J. L., DeGrado, W.F., Influenza A M2 proton channel wild type TM domain bound to S-rimantadine. To be published 2020. DOI: 10.2210/pdb6us8/pdb

32. Knight, C.; Lindberg, G. E.; Voth, G. A., Multiscale reactive molecular dynamics. J Chem Phys 2012, 137 (22), 22A525. DOI: 10.1063/1.4743958

33. Yamashita, T.; Peng, Y.; Knight, C.; Voth, G. A., Computationally Efficient Multiconfigurational Reactive Molecular Dynamics. J Chem Theory Comput 2012, 8 (12), 4863–4875. DOI: 10.1021/ct3006437

34. Nelson, J. G.; Peng, Y.; Silverstein, D. W.; Swanson, J. M., Multiscale Reactive Molecular Dynamics for Absolute pKa Predictions and Amino Acid Deprotonation. J Chem Theory Comput 2014, 10 (7), 2729–2737. DOI: 10.1021/ct500250f

35. Lee, S.; Liang, R.; Voth, G. A.; Swanson, J. M., Computationally Efficient Multiscale Reactive Molecular Dynamics to Describe Amino Acid Deprotonation in Proteins. J Chem Theory Comput 2016, 12 (2), 879–91. DOI: 10.1021/acs.jctc.5b01109

36. Biswas, R.; Tse, Y. L.; Tokmakoff, A.; Voth, G. A., Role of Presolvation and Anharmonicity in Aqueous Phase Hydrated Proton Solvation and Transport. J Phys Chem B 2016, 120 (8), 1793–804. DOI: 10.1021/acs.jpcb.5b09466

37. Knight, C.; Voth, G. A., The curious case of the hydrated proton. Acc Chem Res 2012, 45 (1), 101–9. DOI: 10.1021/ar200140h

38. de Grotthuss, C. J. T., Sur la décomposition de l’eau et des corps qu’elle tient en dissolution á l’aide de l’électricité galvanique. Annales de Chimie 1806, LVIII, 54–74.

39. Agmon, N., The Grotthuss Mechanism. Chem. Phys. Lett. 1995, 244, 456–462.

40. Lee, S.; Swanson, J. M.; Voth, G. A., Multiscale Simulations Reveal Key Aspects of the Proton Transport Mechanism in the ClC-ec1 Antiporter. Biophys J 2016, 110 (6), 1334–45. DOI: 10.1016/j.bpj.2016.02.014

41. Liang, R.; Swanson, J. M. J.; Peng, Y.; Wikström, M.; Voth, G. A., Multiscale simulations reveal key features of the proton-pumping mechanism in cytochrome c oxidase. Proceedings of the National Academy of Sciences 2016, 113, 7420–7425.

42. Liang, R.; Swanson, J. M. J.; Wikstrom, M.; Voth, G. A., Understanding the essential proton-pumping kinetic gates and decoupling mutations in cytochrome c oxidase. Proc Natl Acad Sci U S A 2017, 114 (23), 5924–5929. DOI: 10.1073/pnas.1703654114

43. Parker, J. L.; Li, C.; Brinth, A.; Wang, Z.; Vogeley, L.; Solcan, N.; Ledderboge-Vucinic, G.; Swanson, J. M. J.; Caffrey, M.; Voth, G. A.; Newstead, S., Proton movement and coupling in the POT family of peptide transporters. Proc Natl Acad Sci U S A 2017, 114 (50), 13182–13187. DOI: 10.1073/pnas.1710727114

44. Mayes, H. B.; Lee, S.; White, A. D.; Voth, G. A.; Swanson, J. M. J., Multiscale Kinetic Modeling Reveals an Ensemble of Cl(-)/H(+) Exchange Pathways in ClC-ec1 Antiporter. J Am Chem Soc 2018, 140 (5), 1793–1804. DOI: 10.1021/jacs.7b11463

45. Wang, Z.; Swanson, J. M. J.; Voth, G. A., Modulating the Chemical Transport Properties of a Transmembrane Antiporter via Alternative Anion Flux. J Am Chem Soc 2018, 140 (48), 16535–16543. DOI: 10.1021/jacs.8b07614

46. Watkins, L. C.; Liang, R.; Swanson, J. M. J.; DeGrado, W. F.; Voth, G. A., Proton-Induced Conformational and Hydration Dynamics in the Influenza A M2 Channel. J Am Chem Soc 2019, 141 (29), 11667–11676. DOI: 10.1021/jacs.9b05136

47. Li, C.; Yue, Z.; Espinoza-Fonseca, L. M.; Voth, G. A., Multiscale Simulation Reveals Passive Proton Transport Through SERCA on the Microsecond Timescale. Biophys J 2020, 119 (5), 1033–1040. DOI: 10.1016/j.bpj.2020.07.027

48. Wang, Z.; Swanson, J. M. J.; Voth, G. A., Local conformational dynamics regulating transport properties of a Cl(-) /H(+) antiporter. J Comput Chem 2020, 41 (6), 513–519. DOI: 10.1002/jcc.26093

49. Watkins, L. C.; DeGrado, W. F.; Voth, G. A., Influenza A M2 Inhibitor Binding Understood through Mechanisms of Excess Proton Stabilization and Channel Dynamics. J Am Chem Soc 2020, 142 (41), 17425–17433. DOI: 10.1021/jacs.0c06419

50. Humphrey, W.; Dalke, A.; Schulten, K., VMD -- V isual M olecular D ynamics. Journal of Molecular Graphics 1996, 14, 33–38.

51. Thomaston, J. L.; Alfonso-Prieto, M.; Woldeyes, R. A.; Fraser, J. S.; Klein, M. L.; Fiorin, G.; DeGrado, W. F., High-resolution structures of the M2 channel from influenza A virus reveal dynamic pathways for proton stabilization and transduction. Proc Natl Acad Sci U S A 2015, 112 (46), 14260–5. DOI: 10.1073/pnas.1518493112

52. Ma, C.; Polishchuk, A. L.; Ohigashi, Y.; Stouffer, A. L.; Schon, A.; Magavern, E.; Jing, X.; Lear, J. D.; Freire, E.; Lamb, R. A.; DeGrado, W. F.; Pinto, L. H., Identification of the functional core of the influenza A virus A/M2 proton-selective ion channel. Proc Natl Acad Sci U S A 2009, 106 (30), 12283–8. DOI: 10.1073/pnas.0905726106

53. Day, T. J. F.; Soudackov, A. V.; Cuma, M.; Schmitt, U. W.; Voth, G. A., A second generation multistate empirical valence bond model for proton transport in aqueous systems. Journal of Chemical Physics 2002, 117 (12), 5839–5849. DOI: 10.1063/1.1497157

54. Plimpton, S., Fast Parallel Algorithms for Short-Range Molecular Dynamics. Journal of Computational Physics 1995, 117 (1), 1–19. DOI: 10.1006/jcph.1995.1039

55. Sugita, Y.; Kitao, A.; Okamoto, Y., Multidimensional replica-exchange method for free-energy calculations. The Journal of Chemical Physics 2000, 113 (15), 6042–6051. DOI: 10.1063/1.1308516

56. Bonomi, M. B., G.; Camilloni, C.; et al., Promoting transparency and reproducibility in enhanced molecular simulations. Nat Methods 2019, 16 (8), 670–673. DOI: 10.1038/s41592-019-0506-8

57. Tribello, G. A.; Bonomi, M.; Branduardi, D.; Camilloni, C.; Bussi, G., PLUMED 2: New feathers for an old bird. Computer Physics Communications 2014, 185 (2), 604–613. DOI: 10.1016/j.cpc.2013.09.018

58. Konig, P. H.; Ghosh, N.; Hoffmann, M.; Elstner, M.; Tajkhorshid, E.; Frauenheim, T.; Cui, Q., Toward theoretical analysis of long-range proton transfer kinetics in biomolecular pumps. J Phys Chem A 2006, 110 (2), 548–63. DOI: 10.1021/jp052328q

59. Riccardi, D.; Konig, P.; Prat-Resina, X.; Yu, H.; Elstner, M.; Frauenheim, T.; Cui, Q., “Proton holes” in long-range proton transfer reactions in solution and enzymes: A theoretical analysis. J Am Chem Soc 2006, 128 (50), 16302–11. DOI: 10.1021/ja065451j

60. Liang, R.; Swanson, J. M.; Voth, G. A., Benchmark Study of the SCC-DFTB Approach for a Biomolecular Proton Channel. J Chem Theory Comput 2014, 10 (1), 451–462. DOI: 10.1021/ct400832r

61. Lee, C.; Yang, W.; Parr, R. G., Development of the Colle-Salvetti correlation-energy formula into a functional of the electron density. Phys Rev B Condens Matter 1988, 37 (2), 785–789. DOI: 10.1103/physrevb.37.785

62. Becke, A. D., Density-functional exchange-energy approximation with correct asymptotic behavior. Phys Rev A Gen Phys 1988, 38 (6), 3098–3100. DOI: 10.1103/physreva.38.3098

63. Grimme, S.; Antony, J.; Ehrlich, S.; Krieg, H., A consistent and accurate ab initio parametrization of density functional dispersion correction (DFT-D) for the 94 elements H-Pu. J Chem Phys 2010, 132 (15), 154104. DOI: 10.1063/1.3382344

64. Lippert, B. G.; Parrinello, J. H.; Michele, A hybrid Gaussian and plane wave density functional scheme. Molecular Physics 2010, 92 (3), 477–488. DOI: 10.1080/002689797170220

65. Hartwigsen, C.; Goedecker, S.; Hutter, J., Relativistic separable dual-space Gaussian pseudopotentials from H to Rn. Physical Review B 1998, 58 (7), 3641–3662. DOI: 10.1103/PhysRevB.58.3641

66. VandeVondele, J.; Krack, M.; Mohamed, F.; Parrinello, M.; Chassaing, T.; Hutter, J., Quickstep: Fast and accurate density functional calculations using a mixed Gaussian and plane waves approach. Computer Physics Communications 2005, 167 (2), 103–128. DOI: 10.1016/j.cpc.2004.12.014

67. Wu, H.; Mey, A. S.; Rosta, E.; Noe, F., Statistically optimal analysis of state-discretized trajectory data from multiple thermodynamic states. J Chem Phys 2014, 141 (21), 214106. DOI: 10.1063/1.4902240

68. Scherer, M. K.; Trendelkamp-Schroer, B.; Paul, F.; Perez-Hernandez, G.; Hoffmann, M.; Plattner, N.; Wehmeyer, C.; Prinz, J. H.; Noe, F., PyEMMA 2: A Software Package for Estimation, Validation, and Analysis of Markov Models. J Chem Theory Comput 2015, 11 (11), 5525–42. DOI: 10.1021/acs.jctc.5b00743

69. Zhou, H. X., A theory for the proton transport of the influenza virus M2 protein: extensive test against conductance data. Biophys J 2011, 100 (4), 912–21. DOI: 10.1016/j.bpj.2011.01.002

