## Supplemental File for "Multiscale Simulation of an Influenza A M2 Channel Mutant Reveals Key Features of Its Markedly Different Proton Transport Behavior"

### QM/MM Center of Excess Charge (CEC) Definition

The CEC coordinate in the QM/MM simulations captures the delocalized nature of the excess proton, and is defined as[1]

$$\vec{\xi} = \sum_{i=1}^{N_H} \vec{r}^{H_i} - \sum_{j=1}^{N_X} w^{X_j}\vec{r}^{X_j} - \sum_{i=1}^{N_H}\sum_{j=1}^{N_X} f_{SW}\left(d_{X_jH_i}\right)\left(\vec{r}^{H_i} - \vec{r}^{X_j}\right) + \vec{\xi}_{correct} \qquad (1)$$

where $X_j$'s are the histidine nitrogen atoms and water oxygen atoms in the QM region, and $H_i$'s are the hydrogen atoms bound to those heavy atoms in the QM region. The $w^{X_j}$'s are the hydrogen coordination numbers of the heavy atoms in its molecule's least protonated state during the PT process—for water oxygen atoms this value is 2 (two hydrogens per oxygen), and for His37 nitrogen atoms it is 0.50 in the +0 state (four hydrogens shared by eight His37 nitrogen atoms), 0.625 in the +1 state, and 0.75 in the +2 state. The $d_{X_jH_i}$ variable is the distance between atoms $X_j$ and $H_i$. The function $f_{SW}\left(d_{X_jH_i}\right)$ measures the current coordination number of $H_i$ to $X_j$: $f_{SW}\left(d_{X_jH_i}\right) = 1/1 + \exp\left[(d_{X_jH_i} - r_{SW})/d_{SW}\right]$. The parameters are set to $d_{SW} = 0.04$ Å and $r_{SW} = 1.25$ Å.[2]

The correction term $\vec{\xi}_{correct}$ was previously introduced to correct for the contribution due to the presence of multiple protons around multiple protonatable sites in the His37 tetrad[3]

$$\vec{\xi}_{correct} = \frac{1}{8}\sum_{i=1}^{8}\sum_{j=1}^{8} m_i\left(\vec{r}^{X_j} - \vec{r}^{X_i}\right) \qquad (2)$$

where X denotes one of the eight nitrogen atoms of the His37 tetrad, and $m_i$ switches from 1 to 0 as nitrogen atom $i$ is deprotonated:

$$m_i = \sum_{H_j \in \{H\}} f_{SW}\left(d_{X_i,H_j}\right)^{16} \Bigg/ \sum_{H_j \in \{H\}} f_{SW}\left(d_{X_i,H_j}\right)^{15} . \qquad (3)$$

This formulation has been shown to adequately describe the excess proton position in QM/MM simulations of biological PT channels[1, 2, 4] and has been used in our previous work with M2.[3, 5]

**Supplemental References**

1. Konig, P. H.; Ghosh, N.; Hoffmann, M.; Elstner, M.; Tajkhorshid, E.; Frauenheim, T.; Cui, Q., Toward theoretical analysis of long-range proton transfer kinetics in biomolecular pumps. *J Phys Chem A* **2006,** *110* (2), 548-63.
2. Riccardi, D.; Konig, P.; Prat-Resina, X.; Yu, H.; Elstner, M.; Frauenheim, T.; Cui, Q., "Proton holes" in long-range proton transfer reactions in solution and enzymes: A theoretical analysis. *J Am Chem Soc* **2006,** *128* (50), 16302-11.
3. Liang, R.; Li, H.; Swanson, J. M.; Voth, G. A., Multiscale simulation reveals a multifaceted mechanism of proton permeation through the influenza A M2 proton channel. *Proc Natl Acad Sci U S A* **2014,** *111* (26), 9396-401.
4. Liang, R.; Swanson, J. M.; Voth, G. A., Benchmark Study of the SCC-DFTB Approach for a Biomolecular Proton Channel. *J Chem Theory Comput* **2014,** *10* (1), 451-462.
5. Liang, R.; Swanson, J. M. J.; Madsen, J. J.; Hong, M.; DeGrado, W. F.; Voth, G. A., Acid activation mechanism of the influenza A M2 proton channel. *Proc Natl Acad Sci U S A* **2016,** *113* (45), E6955-E6964.
